# DiSiR: a software framework to identify ligand-receptor interactions at subunit level from single-cell RNA-sequencing data

**DOI:** 10.1101/2022.03.25.485741

**Authors:** Milad R. Vahid, Andre Kurlovs, Franck Auge, Reza Olfati-Saber, Emanuele de Rinaldis, Franck Rapaport, Virginia Savova

**Affiliations:** Artificial Intelligence & Deep Analytics (AIDA), Data & Data Science (DDS), Sanofi R&D, 50 Binney Street, Cambridge, MA 02142, USA; Immunology and Inflammation Research Therapeutic Area, Sanofi, 270 Albany Street, Cambridge, MA 02139, USA; Artificial Intelligence & Deep Analytics (AIDA) Group, Data & Data Science (DDS), 1, Av Pierre Brossolette 91385, Chilly-Mazarin, France

## Abstract

Most of cell-cell interactions and crosstalks are mediated by ligand-receptor interactions. The advent of single-cell RNA-sequencing (scRNA-seq) techniques has enabled characterizing tissue heterogeneity at single-cell level. Over the past recent years, several methods have been developed to study ligand-receptor interactions at cell type level using scRNA-seq data. However, there is still no easy way to query the activity of a specific user-defined signaling pathway in a targeted way or map the interactions of the same subunit with different ligands as part of different receptor complexes. Here, we present DiSiR, a fast and easy-to-use permutation-based software framework to investigate how individual cells are interacting with each other by analyzing signaling pathways of multi-subunit ligand-activated receptors from scRNA-seq data, not only for available curated databases of ligand-receptor interactions, but also for interactions that are not listed in these databases. We show that, when utilized to infer melanoma disease map on a gold-standard dataset, DiSiR outperforms other well-known permutation-based methods, e.g., CellPhoneDB and ICELLNET. To demonstrate DiSiR’s utility in exploring data and generating biologically relevant hypotheses, we apply it to COVID lung and rheumatoid arthritis (RA) synovium scRNA-seq data and highlight potential differences between inflammatory pathways at cell type level for control vs. disease samples.

## Introduction

Ligand-receptor interactions are crucial for the development and normal function of multicellular organisms. It comes as no surprise that drugs commonly produce their therapeutic effects by binding to specific receptors located on specific types of cells. Therefore, dissecting signaling pathways of the ligand-activated receptors at cell type level, is vital to further understanding how multicellular organisms function, and to aid in deciphering drugs’ mechanisms of action and engineering future pharmaceuticals.

The emergence of novel single-cell RNA-sequencing (scRNA-seq) techniques made it possible to study cellular compositions in normal tissues and even more heterogeneous tissues, such as cancerous tissues. Most available single-cell data analysis methods only focus on the prediction of cell type labels and cellular differentiation trajectories [1-3]. However, it is now possible to measure the gene expressions of ligands and receptors in different cell types and systematically decode intercellular communication networks at cell level and their alterations in different diseases.

Over the past recent years, many computational tools have been developed to study ligand-receptor (LR) interactions at cell type level using scRNA-seq data. Armingol et al. [4] have categorized available computational tools, based on their underlying mathematical/statistical model, into the following main categories: (1) differential combination (or differentially expressed genes)-based methods [5-8], (2) expression permutation-based methods [1, 2, 9-11], (3) graph (or network)-based methods [12-15], and (4) array (or tensor)-based methods [16].

A well-known example of the first group is PyMINEr [7] that identifies all non-parametric Spearman’s correlations between genes, creating a graph network in which a connection between two genes indicates that their expression is correlated. Then, the integration of the graph structure and cell type enrichment analyses enables the identification of cell type-specific expression patterns. In SingleCellSignalR [9], as an example of the methods which belong to the second group, in the first step, the contents of existing databases that contain LR pairs, according to literature and experimental data, are compiled. The compiled database, referred to as LRdb, which includes 3251 LR interactions, was built from FANTOM5 [17], HPRD [18], HPMR [19], the IUPHAR/BPS Guide to Pharmacology [20], and UniProtKB/Swissprot [21]. Then, the validity of each potential LR pair according to LRdb is estimated by the computation of a score, called the regularized product score, based on gene expression in the respective cell types. Efremova et al. generalized permutation-based approaches for the cases of LR complexes with multi-subunits and introduced CellPhoneDB [22, 23] as a powerful approach to identify cellular communications in scRNA-seq data. ICELLNET [10] is another example of permutation-based methods that also accounts for multiple LR subunits and is based on multiplying the geometric means of ligand and receptor expressions. In NicheNet [12], which can be classified as a graph-based method, LR links between interacting cells are obtained based on a score calculated by comparing gene expression data with the ligand-signaling network using Personalized PageRank algorithm [24]. As an important effort to use linear algebraic and tensor-based approaches to better understand cell signaling networks, Tsuyuzaki et al. have recently introduced scTensor using non-negative tensor decomposition [16]. In this method, in contrast to conventional methods, cell-cell interactions (CCIs) are regarded as hypergraphs between graphs made of cell types (nodes) and ligand-receptor pairs (edges) and the corresponding data structure is considered as a tensor, not a two-dimensional matrix. Non-negative Tucker decomposition [25] is then used to extract CCIs from the CCI-tensor.

These methods suffer from a main drawback as they are not designed to focus on specific individual LR interactions, especially in the case that both ligands and receptors contain multiple subunits, and instead also interrogate all the interactions that are listed in their built-in database. In this paper, we present a fast and easy-to-use computational method, referred to as DiSiR (DiSiR: <u>Di</u>mer <u>Si</u>gnal <u>R</u>eceptor Analysis), which builds upon the framework from existing expression permutation-based methods such as CellPhoneDB by addressing this drawback. Our method accepts user-defined LR input, including readily available curated LR databases, and has been specifically optimized to analyze complex pathways involving multi-subunit receptor complexes with multi-subunit ligands, or hypothetical interactions. Unlike other methods which emphasize overall number of interactions across cell types, we have focused on visualizing which cell types interact directionally via the specific signaling mechanism of interest. Here, we evaluate the performance of DiSiR using a “gold-standard” LR interaction database [17] from the literature. Zhou et al. [26] analyzed these interactions in scRNA-seq data from melanoma patients and generated a cell-cell communication network for melanoma. We use this network as the ground truth and benchmark the performance of DiSiR against well-known available permutation-based methods, e.g., CellPhoneDB [22] and ICELLNET [10].

## Results and Discussion

DiSiR uses single-cell gene expression matrix, cell type annotations, and a list of desired ligand-receptor interactions at subunit level as input (Fig 1). DiSiR identifies ligand “L”–receptor “R” interaction between cell types “A” and “B” based on the products of normalized expressions of “L” subunits by cell type “A” and normalized expressions of “R” subunits by cell type “B”. This interaction is considered by DiSiR as significant if these products are significantly higher than the result of random shuffling of cell type labels for all “L–R” subunit combinations. We filter out interactions between cell types with low fractions of cells expressing the ligand/receptor subunits (see “Materials and Methods” section).

**Fig 1.**
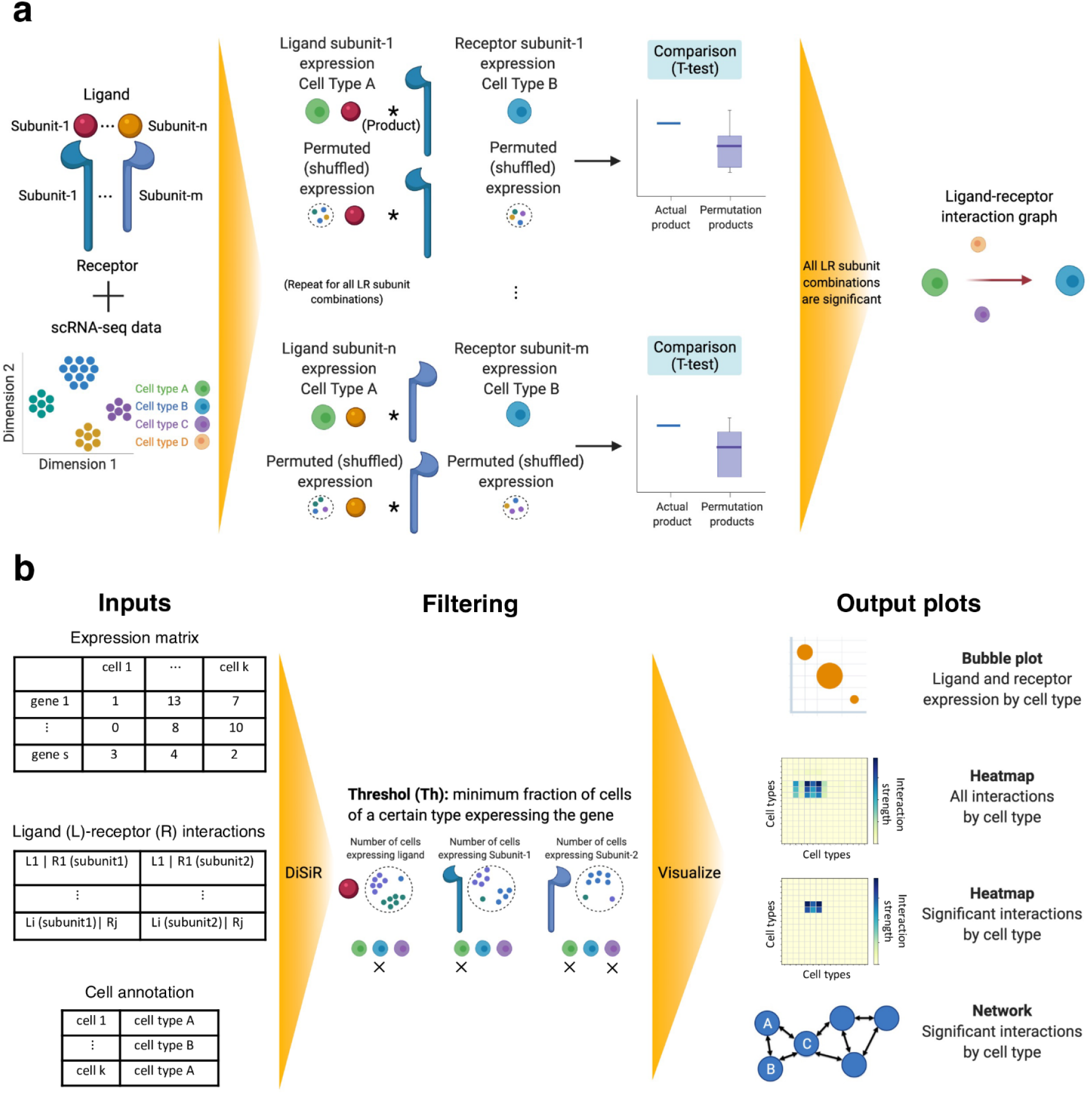
Schematic overview and illustration of the ease-of-use of DiSiR framework. **(a)** DiSiR uses 1) single-cell gene expression matrix, 2) cell type annotations, and 3) list of desired ligand-receptor interactions at subunit level as input data. DiSiR then characterizes the ligand “L”–receptor “R” interaction between cell types “A” and “B” based on the products of expressions of “L” subunits by cell type “A” and expressions of “R” subunits by cell type “B”. This interaction is identified by DiSiR as a significant interaction if these products are significantly higher than random shuffling results for all “L–R” subunit combinations. **(b)** Diagram that shows the workflow of data processing, filtering, and visualization of the obtained results in DiSiR in which users only need to input the three datasets mentioned in part (a). Weak interactions are then filtered out if the fraction of cells expressing the ligand (or receptor) within its corresponding cell type is less than a threshold *Th*. DiSiR also provides an interactive graph-based representation module and heatmap/bubble plots to show the resulting interactions for complex signaling pathways at different levels.

### Analysis of individual multi-subunit ligand-receptor interactions at cell type level

Here, we have applied DiSiR on the RA scRNA-seq data to analyze the S100A8/A9 (calprotectin) signaling pathway (Fig 2). We have shown the result for the optimal threshold in Fig 2 and included the results across different threshold levels in S1 Fig. The SPRING plot of the RA synovium revealed a total of 8,990 cells (Fig 2a). Of those cells, all but 691 were annotated by SignacX. To test DiSiR, we chose the calprotectin-toll-like-receptor 4 (CLP/TLR4) pathway, in which the proteins encoded by S100A8 and S100A9 combine into a dimer and bind TLR4. This pathway is a known biomarker of RA [27] that is currently absent from most available databases, including the CellPhoneDB database. This result serves as proof of concept that DiSiR is capable of quickly interrogating a pathway, as well as sheds additional light on the biology of this interaction. The higher weights of the links connected to macrophages (Fig. 2c) suggest that macrophages are the primary signaling cells, signaling primarily to themselves and to monocytes. Note that we use the following notation in the interaction graph: there is a link L1 | R1 + L2 | R2 between two nodes C1 and C2, if cell types C1 and C2 communicate with each other through the ligand L-receptor R interactions, where L1, L2 and R1, R2 are subunits of L and R, respectively. In addition, some signaling took place in cells that are not part of the immune system (annotated as ‘Fibroblasts’ and ‘NonImmune’). These findings are consistent with the existing paradigm [28] that CLP is released by phagocytes and TLR4 binds to it in phagocytic cells as well as in cells that are not immune, but also contains the resolution and the granularity that is more specific to the AMP RA Phase1 dataset and that can help narrow down the exact mechanism of CLP signaling.

**Fig 2.**
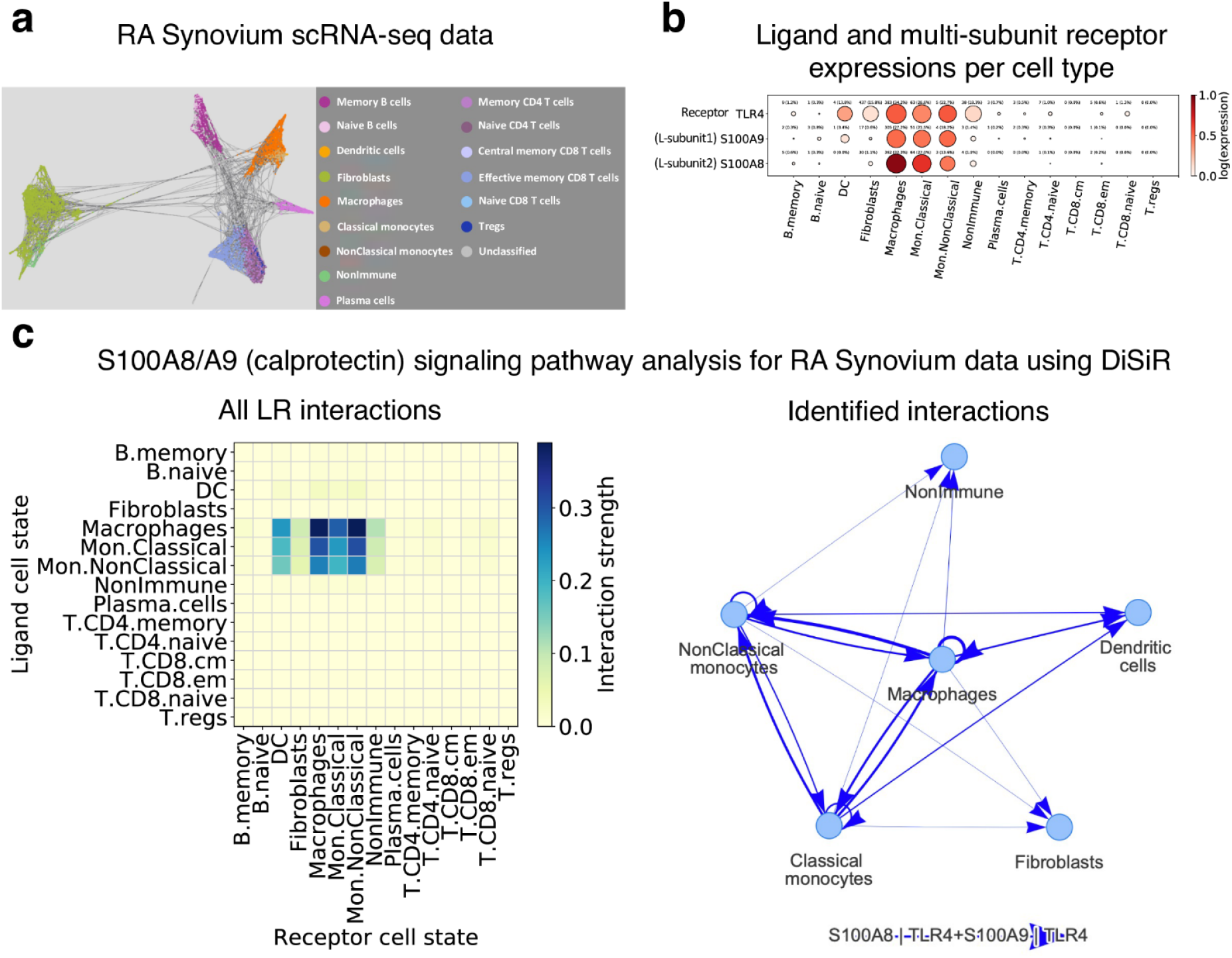
Analysis of Calprotectin signaling pathways at cell type level from RA Synovium scRNA-seq data. **(a)** UMAP representation of RA Synovium scRNA-seq data and the corresponding cell type label assigned to each cell. **(b)** Bubble plot illustrating max-normalized average expressions of calprotectin signaling pathway components including ligand subunits, S100A8 and S100A9, and receptor TLR4 per cell type (color of the circles) and fraction of cells expressing them within its corresponding cell type (size of the circles). **(c)** Left panel is the heatmap depicting all cell-cell communication through S100A8|TLR4 + S100A9|TLR4 interactions (when both interactions are presented). The colormap shows the strength of interaction between two cell types. Right panel shows the significant S100A8-TLR4 and S100A9-TLR4 interaction graphs between different cell types identified by DiSiR. The thickness of the graph edges corresponds to the interaction strength.

In addition to CLP, we also tested DiSiR on another common pathway, the IL6 classical signaling pathway that requires both IL6ST and IL6R receptor subunits (S2 Fig). DiSiR has predicted that in RA synovium, IL6 is primarily secreted by B-cells, while CD4+ T-cells are the main immune cell types on the receiving end. As with CLP, the IL6 pathway is also elevated in RA. Consistent with the recent review of the role of IL6 in RA [29], B-cells are major secretors of IL6. However, another immune cell type associated with IL6 secretion in the review – macrophages – were not major secretors of IL6 in the AMP RA dataset. We did not observe a high level of expression of IL6 in the macrophages from the AMP RA dataset (see S2b Fig) to account for the macrophage discrepancy. The role of IL6 signaling to CD4+ T-cells has also attracted researchers’ attention [30].

### Validation and benchmarking analysis of DiSiR across different methods

In order to evaluate the performance of DiSiR, we use a LR interaction database [17] from the literature. This database is a part of FANTOM5 and includes a table reporting the list of ∼2500 LR pairs over 144 primary cell types, which represents a large-scale map of cell-to-cell communication in human. Zhou et al. [26] investigated these interactions in the transcriptomes of ∼4,000 single cells isolated from melanoma patients, including both malignant tumor cells and stromal and immune cells. By systematically comparing the LR pair expression patterns among seven major cell types present in the melanoma-derived single-cell data (melanoma cells, T cells, B cells, macrophages, NK cells, cancer-associated fibroblasts (CAF) and endothelial cells), the authors were able to build a cell-cell communication map indicating which cell type interacts with each other, based on specific LR links. We use this melanoma disease map as the ground truth and benchmark the performance of DiSiR against well-known available permutation-based methods, e.g., CellPhoneDB and ICELLNET, which are comparable to DiSiR in terms of the required input data, underlying approach, and output results (the melanoma scRNA-seq data that we have used for this analysis is from [31]).

We compute recall (sensitivity), precision, and specificity measures (see the “ROC and precision-recall curves” section for more details) of identifying LR interactions at cell type level, between cancer cells and the other cell types, for different threshold values and use these values to generate the corresponding ROC (Fig 3) and precision-recall curves (S3 Fig) for all methods. The area under the curve (AUC) and calculated sensitivity and specificity scores show that DiSiR outperforms the other methods across different threshold values ranging from 0 and 1.

**Fig 3.**
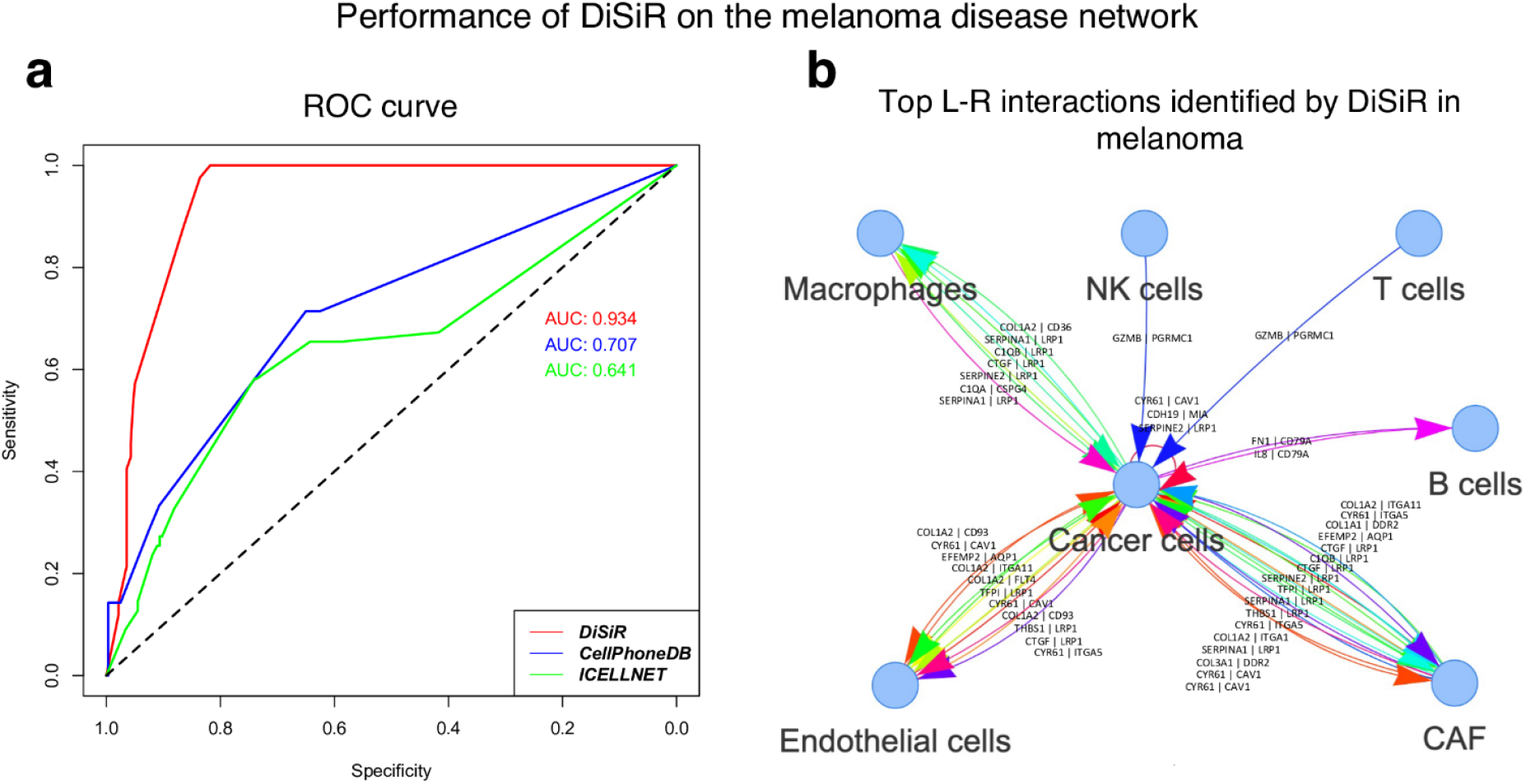
Technical assessment of DiSiR using a gold-standard melanoma cellular network. **(a)** Sensitivity and specificity, across different threshold values, are defined in relation to positive and negative groups for DiSiR vs. CellPhoneDB and ICELLNET (AUC, area under the curve). **(b)** Graph representation of top ligand-receptor interactions between cancer cells and different cell types identified by DiSiR.

We next calculate optimal cut-off points in ROC and precision-recall curves to select the best value for the threshold parameter *Th* using Youden’s J statistic and sensitivity score, respectively. Since the sensitivity criterion is more important for us, we proceed with the optimal threshold obtained in this case (*Th* = 0). This low optimal threshold value suggests that DiSiR performs well even when there are fewer cells expressing the ligands/receptors of interest. We propose to set this optimal value as the default value of the threshold parameter in our tool and use it to analyze other scRNA-seq data sets in this paper, e.g., RA and COVID lung scRNA-seq data.

We have also compared the running time of DiSiR and CellPhoneDB (searching over its entire database) applied to the AMP RA Phase 1 data on our system (2.4 GHz 8-Core Intel Core i9 processor 16GB RAM 2667 MHz DDR4). The execution time of about one minute for DiSiR vs. ∼ 55 minutes for CellPhoneDB, suggests that using DiSiR, it is easy and time efficient to study single interactions between individual components of complex pathways.

### Application of DiSiR to analyze and visualize complex immunoinflammatory signaling pathways in COVID-19 scRNA-seq data

As another application, we studied differences in cell-cell communications through more complex inflammatory signaling pathways for disease versus control samples. For this purpose, we have applied DiSiR on the lung scRNA-seq data to analyze the IL6/IL11 (Fig 4) and IL1RAP (Figs. 5) signaling pathways for COVID-19 vs. healthy samples. As a member of IL6 family cytokines, IL11 competes with IL6 for its joint receptor subunit IL6ST. In fact, IL11 binds to another receptor subunit IL11R and forms a complex with IL6ST which activates the downstream signaling through the activation STAT3, ERK or PI3K pathways [32]. We analyzed these two pathways jointly since they have a shared subunit (Fig 4). Our results depict the potential enriched ligand-receptor interactions between monocytes/macrophages with each other and also, with T cells (CD4, CD8 and Tregs) in the disease stage compared to healthy controls, which is consistent with the results shown in [33] and for COVID patients in a severe stage.

**Fig 4.**
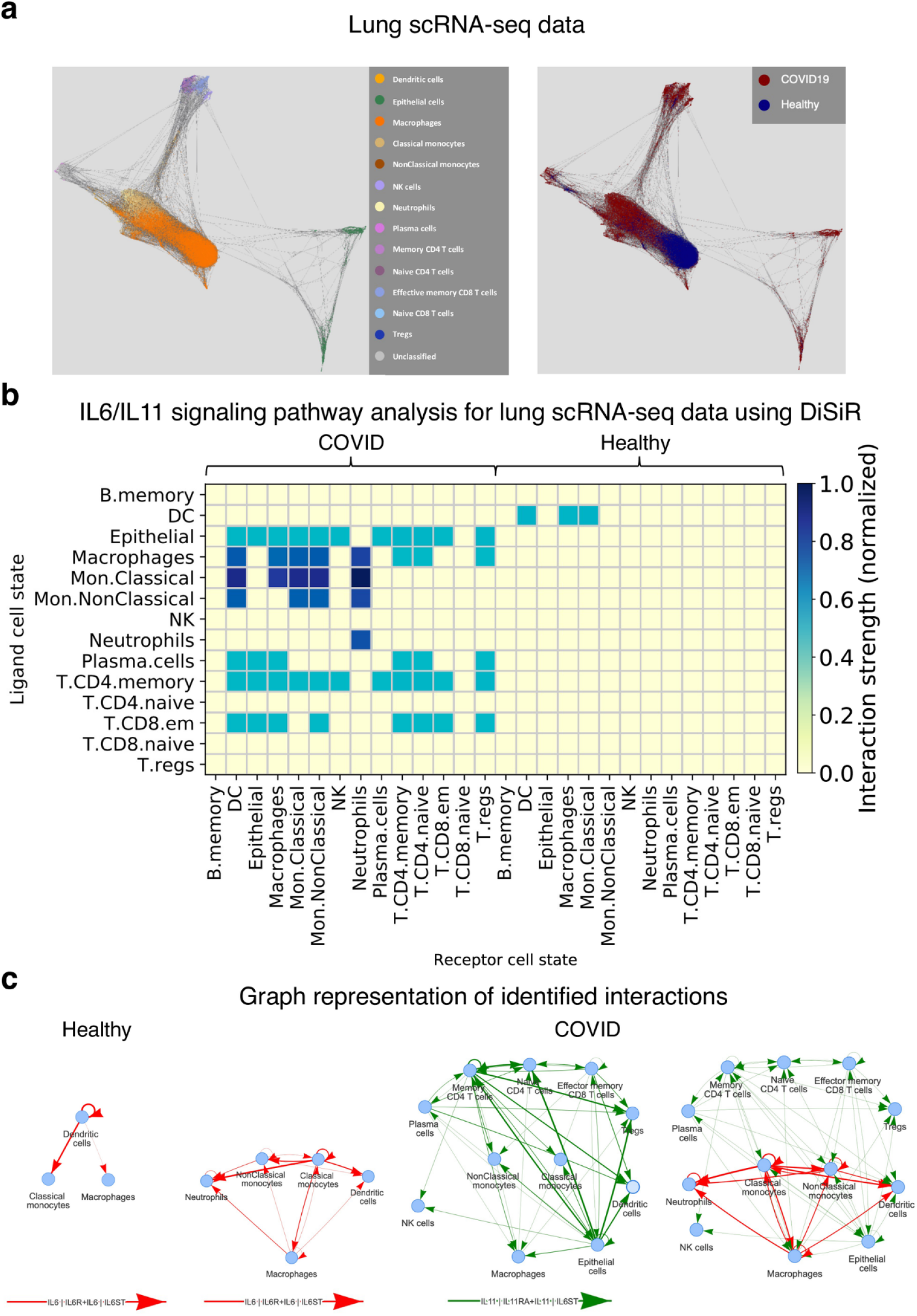
Study of IL6/IL11 signaling pathways for COVID vs. healthy control samples using DiSiR from lung scRNA-seq data. **(a)** UMAP representation of lung scRNA-seq data and corresponding cell type (left] and disease labels e.g., COVID19 or healthy [23], assigned to each cell. **(b)** Heatmap plot illustrating the difference between cell-cell communications identified by DiSiR for the IL6/IL11 signaling pathway between COVID vs. control samples in lung scRNA-seq data. The colormap shows the strength of interaction between two cell types. **(c)** Graph representation of the significant ligand-receptor interactions, involved in the IL6/IL11 signaling pathway, i.e., IL6|IL6R + IL6|IL6ST and IL11|IL116A + IL11|IL6ST, between different cell types identified by DiSiR for healthy vs. COVID samples using the default threshold level, which is equal to 0. The thickness of the graph edges corresponds to the interaction strength.

**Fig 5.**
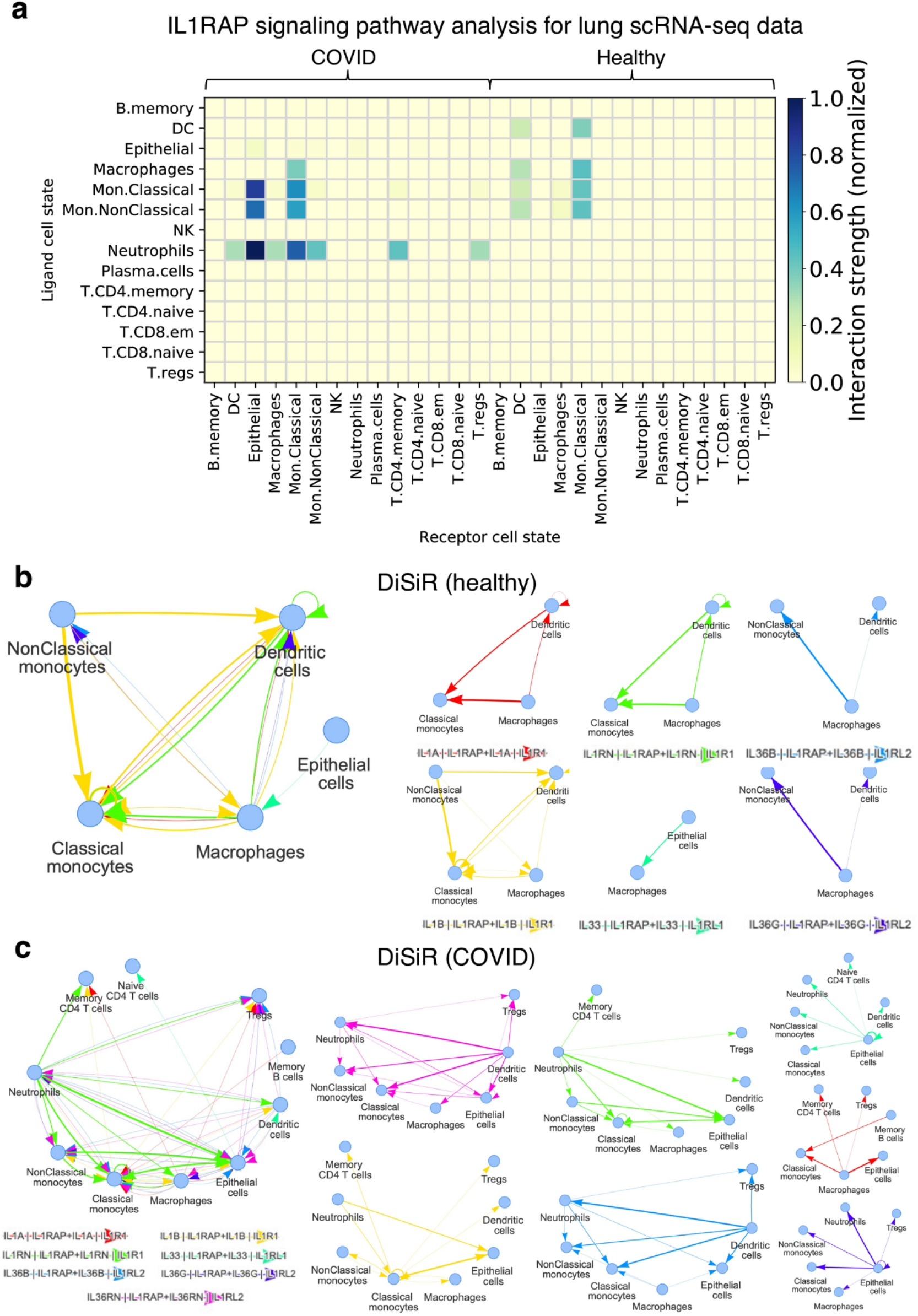
Study and illustrations of complex multi-subunits IL1RAP signaling pathways for COVID vs. control samples using DiSiR from lung scRNA-seq data. **(a)** Heatmap plot illustrating the differences between cell-cell communications identified by DiSiR for IL1RAP signaling pathway between COVID vs. control samples in lung scRNA-seq data. The colormap shows the strength of interaction between two cell types. **(b)** Graph representation of significant ligand-receptor interactions, involved in IL1RAP signaling pathway, e.g., IL1A|IL1RAP + IL1A|IL1R1, IL1B|IL1RAP + IL1B|IL1R1, IL1RN|IL1RAP + IL1RN|IL1RL1, IL33|IL1RAP + IL33|IL1RL1, IL36B|IL1RAP + IL36B|IL1RL2, IL36G|IL1RAP + IL36G|IL1RL2 and IL36RN|IL1RAP + IL36RN|IL1RL2, between different cell types identified by DiSiR for healthy vs. **(c)** COVID samples using the default threshold level, which is equal to 0. The thickness of the graph edges corresponds to the interaction strength.

We also analyzed the IL1RAP pathway which contains more interacting proteins with several shared components. IL1RAP is involved in three signaling pathways that affect many IL1 family cytokines, such as IL1A, IL1B, IL33 and IL36, in different diseases [35]. They then form seven multi-subunit pairs which are illustrated with different colors in Fig 5. It has been shown that IL1RAP and the majority of the components involved in its signaling pathway are differentially expressed between the airways of COVID patients and SARS-cov2 negative patients according to nasopharyngeal swabs data (q-value for IL1RAP, IL1A, IL1B, IL36A and IL36B are 1.97 × 10-^10^, 0.00185, 0.015, 0.05 and 0.023, respectively) [36]. We have applied DiSiR to the COVID data to study IL1RAP signaling pathway at cell type level. As can be seen in Fig 5, we have found enriched LR interactions between ligand-expressing monocytes/macrophages and neutrophils with receptor expressing monocytes/macrophages, epithelial cells, memory CD4 T cells and regulatory T cells (Tregs) in the disease stage.

## Materials and Methods

### Overview of the method

DiSiR is designed to enable identifying single multi-subunit ligand-receptor interactions at cell type level through the following algorithm (Fig 1):

1. Assume the subunits of ligand L and receptor R are denoted by *L*_1_, *L*_2_, …, *L*_*N*_ and *R*_1_, *R*_2_, …, *R*_*M*_, respectively. Also, assume that cells belong to cell types A and B are referred to as clusters A and B.
2. For each *L*_*i*_– *R*_*j*_ interaction, where *K*:={(*L*_*i*_, *R*_*j*_)} ∈ *L*_*n*_, *R*_*m*_)|*n* = 1,…,*N*, = 1,…,*M*} denotes a set of all ligand-receptor subunit pairs listed in the input interactions list, calculate “clusters A and B average expressions” by calculating the average expressions of *L*_*i*_ and *R*_*j*_ over all the cells within clusters A and B, respectively.
3. Generate *k* sets of “shuffled clusters A^’^ and B^’^ average expressions” by randomly permuting the cluster assignment of all cells and re-calculating the average expressions of *L*_*i*_ and *R*_*j*_ over all the cells belonging to the new random clusters A’ and B’ (the default value for *k* is 100).
4. The *L*_*i*_– *R*_*j*_ interaction is considered by DiSiR as a significant interaction if the product of clusters A and B average expressions is significantly higher than the products of shuffled clusters *A*^’^ and *B*^’^ average expressions according to a one-sided paired T-test and multiple hypothesis testing. Multiple hypothesis testing correction is performed using the Benjamini and Hochberg method (results with an adjusted p-value < 0.05 are considered significant).
5. DiSiR identifies a L–R interaction between cell types A and B as significant if *L*_*i*_–*R*_*j*_ interactions are significant for all combinations listed in the input interactions list.

We next explain each step in more details and introduce the filtering approach that we use to remove weak interactions. We also describe the plotting capabilities that we have included in the DiSiR framework in order to better visualize complex signaling pathways.

### Single-cell data sets and data pre-preprocessing steps

We tested DiSiR on a couple of pathways in two publicly available single-cell datasets. We obtained lung COVID-19 single-cell transcriptomic data from GEO145926 [33] and the AMP consortium’s Phase 1 rheumatoid arthritis (RA) data from https://immunogenomics.io/ampra/ [8]. We subsequently filtered the data based on empirically customized standard procedures. We only kept cells for further analysis if they had at least 1000 genes and at most 20% of the genes were mitochondrial (COVID), or if they had at least 200 genes and at most 10% of the genes were mitochondrial (AMP RA Phase 1). We then normalized total barcode counts in each of the two datasets so that each cell had the same total normalized read count. Afterwards, we used SPRING [37], a web-based software tool to visualize high dimensional transcriptomic data, to visualize the data, with 60 (COVID) and 30 principal components (AMP RA Phase 1). We then annotated the cells with SignacX version 2.2.0 [38] – a neural-network-based cell annotation tool that uses expression data and is also aided by the edges from the SPRING plot. Finally, we transferred the resulting annotated expression data into log2 space for DiSiR analysis.

### Filtering approach

DiSiR uses a filtering approach to discard weak interactions that might be associated with noise by thresholding the fraction of cells expressing each ligand/receptor subunit within its corresponding cell type. The user can visualize this metric for combinations of signaling pathway components and cell types using a bubble plot in which the size of each circle is associated with the corresponding fraction and the color of each circle is related to the max-normalized average expression of each ligand/receptor subunit within its corresponding cell type (Fig 2b). A ligand-receptor interaction between two cell types is then retained if this score is higher than a certain threshold level 0 ≤ *Th* ≤ 1 for all ligand/receptor subunits. We provide an analysis framework using precision-recall curves, across all possible threshold values, to determine the optimal value for *Th* (see the “Results and discussion” section as well as S3 Fig). Users also have the freedom to be more stringent on the value of Th in order to focus on stronger interactions.

### Graph and heatmap plot representation

In DiSiR, we visualize output cell-cell interactions in two ways: graph representation and heatmap plots. Graph representation is made of a directed graph in which nodes are associated with the cell types present in the input data set and each edge corresponds to a LR interaction between two cell types, from ligand-expressing cells to receptor-expressing cells (Fig 2c). For a given interaction, if both ligand and receptor are present in the same cell type, then there is a self-loop in the graph on the corresponding node. We use the “visNetwork version 2.1.0” package in R version 4.0.0 with an interactive environment. Heatmap plots illustrate all interactions (including above threshold interactions) between different cell types listed in rows and columns of the heatmaps (Fig 2c). The thickness of links in graph representation and the color intensity in heatmap representation are associated with the strength of interactions between cell types. Single-cell clustering plots were also visualized using SPRING and Harmony [39] version 0.1.0 (for COVID data).

### Ligand-receptor interaction strength

DiSiR identifies significant interactions based on the product of the expressions of ligand and receptor subunits. The L-R interaction strength is defined as

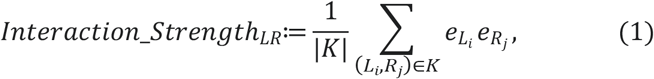

Where 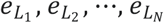 and 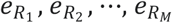, denote log2 TPM expressions of ligand L and receptor R subunits expressed by cell types A and B respectively, *K* = *L*_*i*_, *R*_*j*_)} is a set of all ligand-receptor subunit pairs listed in the input interactions list, and |*K*| is the number of elements of *K*.

### ROC and precision-recall curves

We compare the identified LR interactions with a “gold-standard” LR interaction database at cell type level (see the “Validation and benchmarking analysis of DiSiR across different methods” section). For each cell type, we categorize the identified interactions which are included in the expected interactions as true positives and the ones that are absent from the expected interactions as false positives. The expected interactions that are not matched with any identified interaction are referred to as false negatives. Finally, the interactions that are neither identified by the method nor listed in the expected interactions are true negatives. Denoting the number of true positives by TP, the number of true negatives by TN, the number of false negatives by FN, and the number of false positives by FP, we define the precision (*Pr*), recall or sensitivity (*Sn*), and specificity (*Sp*) measures as

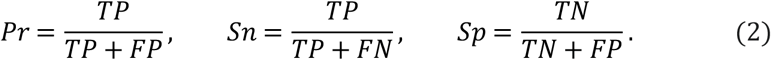

We then use these scores to produce ROC and precision-recall curves and evaluate the performance of DiSiR compared to other similar methods as measured by the area under the curve (AUC).

## Conclusion

In summary, DiSiR is a novel easy-to-use software framework that enables the analysis of complex multi-subunit ligand–multi-subunit receptor interactions from scRNA-seq data, even for the interactions (or genes) that are not listed in available databases. We have applied our method to COVID and RA scRNA-seq data and highlighted potential differences between inflammatory pathways at cell type level for control vs. disease samples.

## Data Availability Statement

All expression transcriptomic data used in this study are publicly available (see the “Single-cell data sets and data pre-preprocessing steps” section). All simulated expression data and the corresponding codes for generating them are available at: https://github.com/miladrafiee/DiSiR. DiSiR V.1.0 was used to generate the results in this work and is freely available for academic research use at the above GitHub page as well.

## Supporting information

**S1 Fig.**
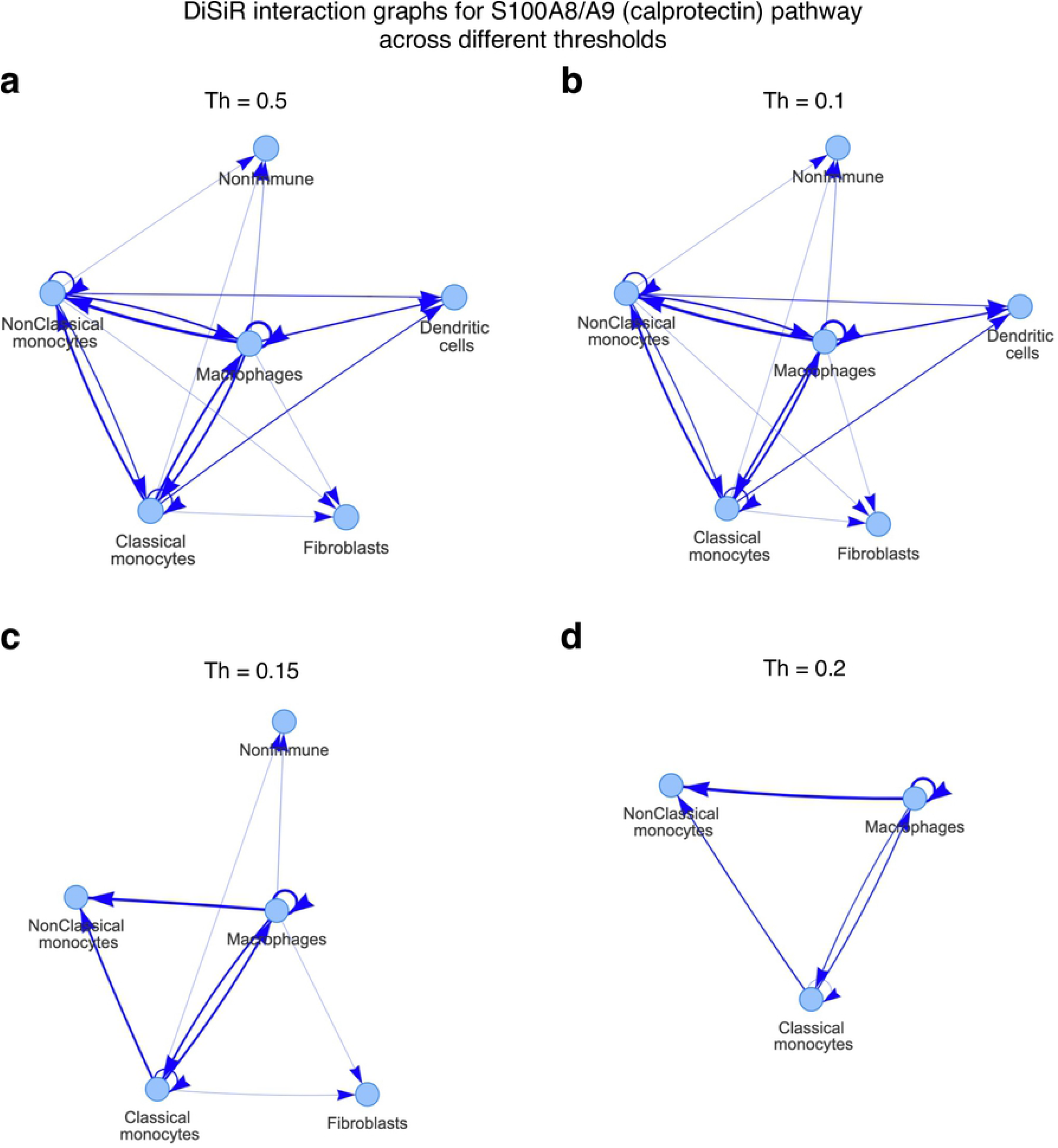
DiSiR interaction graphs for S100A8/A9 (calprotectin) pathway in RA data across different thresholds: **(a)** *Th* = 0.05, **(b)** *Th* = 0.1, **(c)** *Th* = 0.15 and **(d)** *Th* = 0.2.

**S2 Fig.**
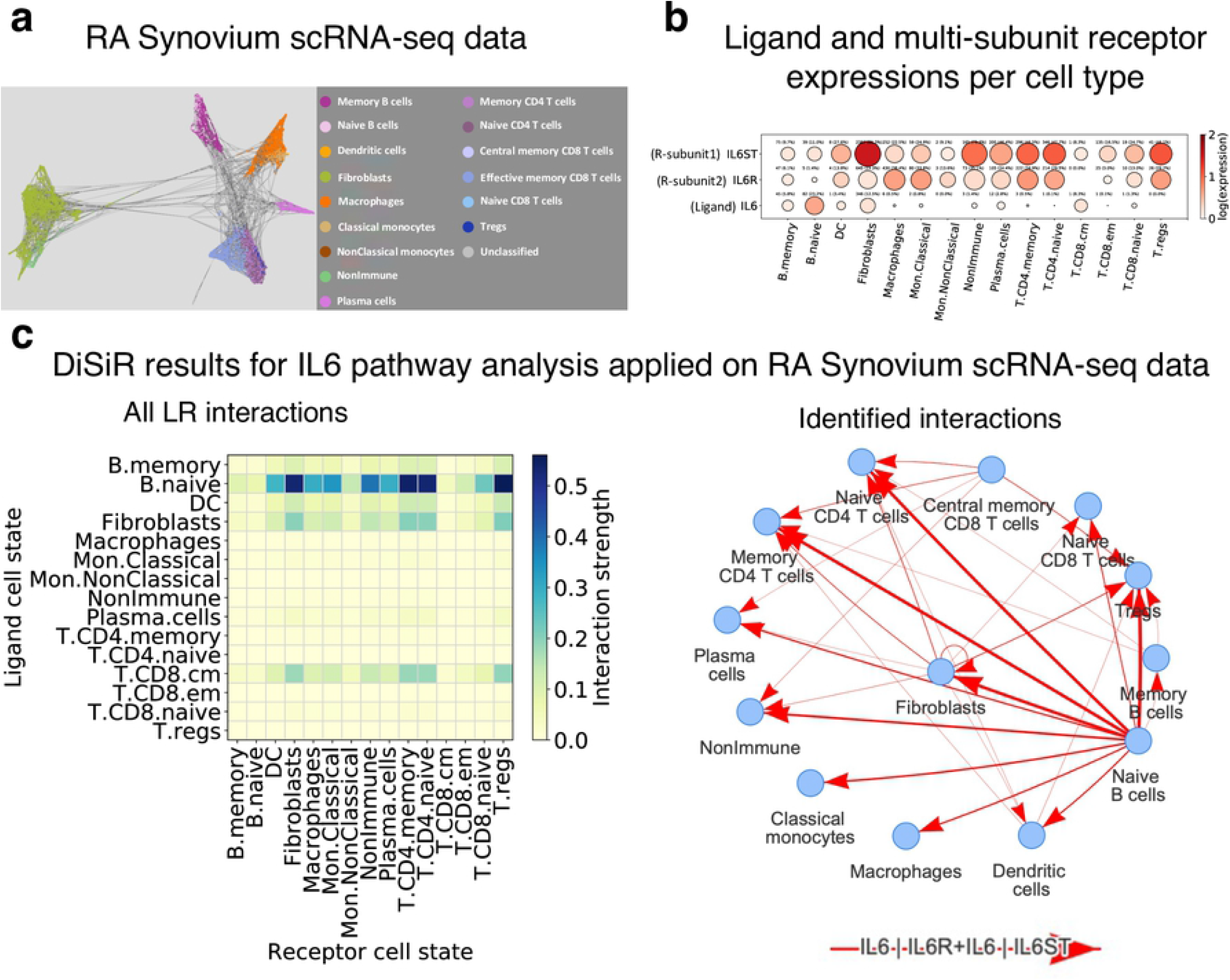
Analysis of IL6 signaling pathways at cell type level from RA Synovium scRNA-seq data. **(a)** UMAP representation of RA Synovium scRNA-seq data and corresponding cell type label assigned to each cell. **(b)** Circle plot illustrating max-normalized average expressions of IL6 signaling pathway components including ligand ILR and receptor subunits, IL6R and IL6ST, per cell type (color of the circles) and fraction of cells expressing them within its corresponding cell type (size of the circles). **(c)** Left figure is the heatmap depicting all cell-cell communication through IL6|IL6R + IL6|IL6ST interactions (when both interactions are presented). The colormap shows the strength of interaction between two cell types. Right graphs show the significant IL6-IL6R and IL6-IL6ST interactions between different cell types identified by DiSiR. The thickness of the graph edges corresponds to the interaction strength.

**S3 Fig.**
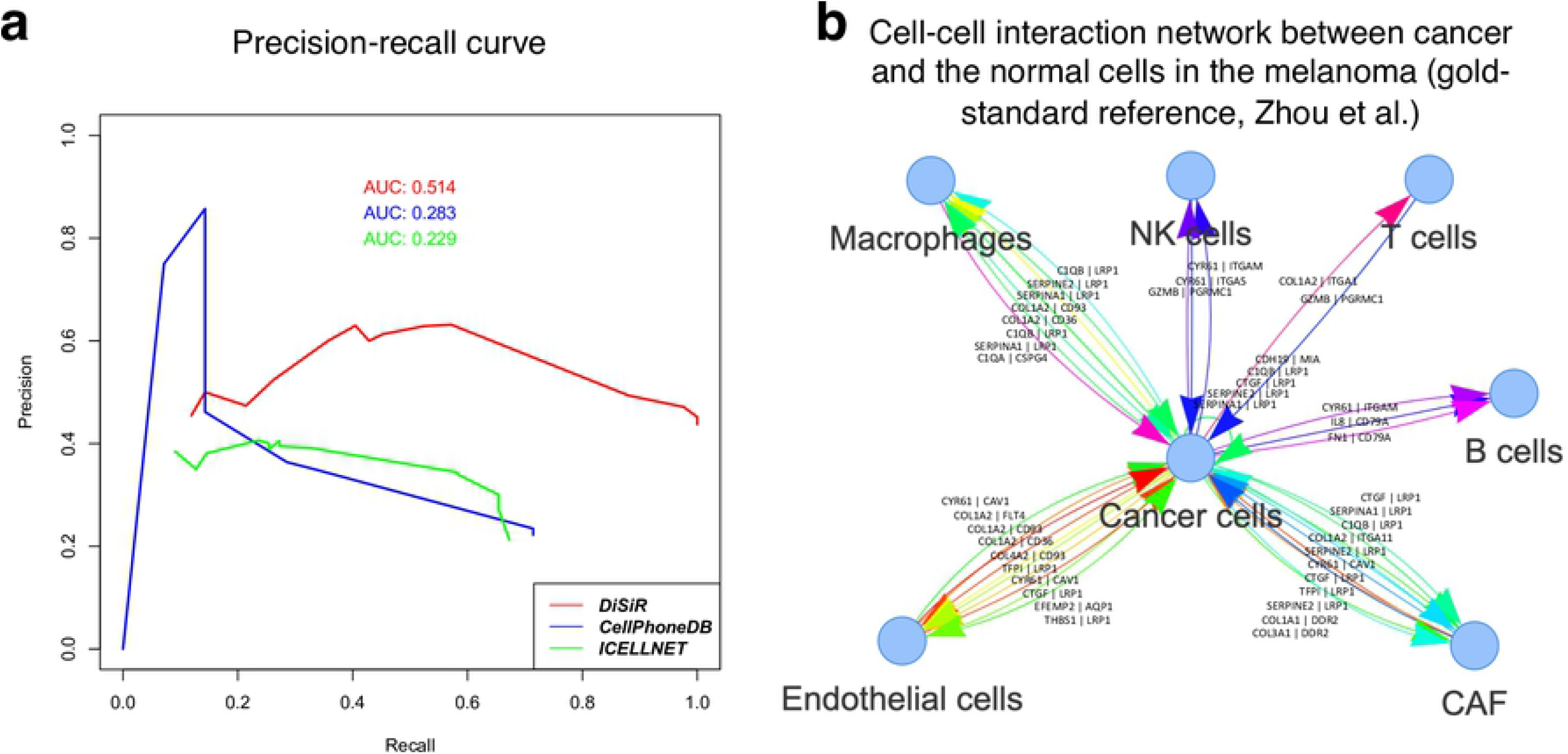
Technical assessment of DiSiR using gold-standard melanoma cellular network based on precision-recall curve. **(a)** Recall and precision, across different threshold values, are defined in relation to positive and negative groups for DiSiR vs. CellPhoneDB and ICELLNET (AUC, area under the curve). Best precision in DiSiR is obtained for *Th* = 0 (optimal threshold value) **(b)** Cell-cell interaction network between cancer and the normal cells in the melanoma (our gold-standard reference) extracted in [26].

## Author Contributions

**Conceptualization:** Virginia Savova, Milad R. Vahid, Franck Auge, Reza Olfati-Saber.

**Data curation:** Milad R. Vahid, Andre Kurlovs, Virginia Savova.

**Formal analysis:** Milad R. Vahid, Andre Kurlovs.

**Funding acquisition:** Franck Auge, Reza Olfati-Saber, Emanuele de Rinaldis, Virginia Savova, Franck Rapaport.

**Investigation:** Milad R. Vahid.

**Methodology:** Milad R. Vahid, Andre Kurlovs, Virginia Savova, Franck Rapaport.

**Project administration:** Virginia Savova, Franck Rapaport.

**Software:** Milad R. Vahid, Andre Kurlovs.

**Validation:** Milad R. Vahid, Andre Kurlovs.

**Visualization:** Milad R. Vahid, Andre Kurlovs.

**Writing – original draft:** Milad R. Vahid, Franck Rapaport.

**Writing – review & editing:** Milad R. Vahid, Andre Kurlovs, Virginia Savova, Franck Rapaport, Franck Auge.

## Acknowledgements

We would like to thank Richa Hanamsagar for insightful discussions.

## References

1. Jin S, Guerrero-Juarez CF, Zhang L, Chang I, Ramos R, Kuan CH, et al. Inference and analysis of cell-cell communication using CellChat. Nature Communications. 2021;12(1):1–20.

2. Stuart T, Butler A, Hoffman P, Hafemeister C, Papalexi E, Mauck WM, 3rd, et al. Comprehensive Integration of Single-Cell Data. Cell. 2019;177(7):1888–902 e21.

3. La Manno G, Soldatov R, Zeisel A, Braun E, Hochgerner H, Petukhov V, et al. RNA velocity of single cells. Nature. 2018;560(7719):494–8.

4. Armingol E, Officer A, Harismendy O, Lewis NE. Deciphering cell-cell interactions and communication from gene expression. Nat Rev Genet. 2021;22(2):71–88.

5. Cillo AR, Kurten CHL, Tabib T, Qi Z, Onkar S, Wang T, et al. Immune Landscape of Viral-and Carcinogen-Driven Head and Neck Cancer. Immunity. 2020;52(1):183–99 e9.

6. Wang Y, Wang R, Zhang S, Song S, Jiang C, Han G, et al. iTALK: an R package to characterize and illustrate intercellular communication. BioRxiv. 2019:p.507871.

7. Tyler SR, Rotti PG, Sun X, Yi Y, Xie W, Winter MC, et al. PyMINEr Finds Gene and Autocrine-Paracrine Networks from Human Islet scRNA-Seq. Cell Rep. 2019;26(7):1951–64 e8.

8. Zhang F, Wei K, Slowikowski K, Fonseka CY, Rao DA, Kelly S, et al. Defining inflammatory cell states in rheumatoid arthritis joint synovial tissues by integrating single-cell transcriptomics and mass cytometry. Nat Immunol. 2019;20(7):928–42.

9. Cabello-Aguilar S, Alame M, Kon-Sun-Tack F, Fau C, Lacroix M, Colinge J. SingleCellSignalR: inference of intercellular networks from single-cell transcriptomics. Nucleic Acids Res. 2020;48(10):e55.

10. Noël F, Massenet-Regad L, Carmi-Levy I, Cappuccio A, Grandclaudon M, Trichot C, et al. Dissection of intercellular communication using the transcriptome-based framework ICELLNET. Nature communications. 2021;12(1):1–16.

11. Dries R, Zhu Q, Dong R, Eng CL, Li H, Liu K, et al. Giotto: a toolbox for integrative analysis and visualization of spatial expression data. Genome Biol. 2021;22(1):78.

12. Browaeys R, Saelens W, Saeys Y. NicheNet: modeling intercellular communication by linking ligands to target genes. Nat Methods. 2020;17(2):159–62.

13. Choi H, Sheng J, Gao D, Li F, Durrans A, Ryu S, et al. Transcriptome analysis of individual stromal cell populations identifies stroma-tumor crosstalk in mouse lung cancer model. Cell Rep. 2015;10(7):1187–201.

14. Cang Z, Nie Q. Inferring spatial and signaling relationships between cells from single cell transcriptomic data. Nature communications. 2020;11(1):1–13.

15. Wang S, Karikomi M, MacLean AL, Nie Q. Cell lineage and communication network inference via optimization for single-cell transcriptomics. Nucleic Acids Res. 2019;47(11):e66.

16. Tsuyuzaki K, Ishii M, Nikaido I. Uncovering hypergraphs of cell-cell interaction from single cell RNA-sequencing data. bioRxiv. 2019:p.566182.

17. Ramilowski JA, Goldberg T, Harshbarger J, Kloppmann E, Lizio M, Satagopam VP, et al. A draft network of ligand-receptor-mediated multicellular signalling in human. Nat Commun. 2015;6(1):7866.

18. Prasad TK, Kandasamy K, Pandey A. Human Protein Reference Database and Human Proteinpedia as discovery tools for systems biology. Methods Mol Biol. 2009;577:67–79.

19. Ben-Shlomo I, Yu Hsu S, Rauch R, Kowalski HW, Hsueh AJ. Signaling receptome: a genomic and evolutionary perspective of plasma membrane receptors involved in signal transduction. Sci STKE. 2003;2003(187):RE9.

20. Harding SD, Sharman JL, Faccenda E, Southan C, Pawson AJ, Ireland S, et al. The IUPHAR/BPS Guide to PHARMACOLOGY in 2018: updates and expansion to encompass the new guide to IMMUNOPHARMACOLOGY. Nucleic Acids Res. 2018;46(D1):D1091–D106.

21. Wu CH, Apweiler R, Bairoch A, Natale DA, Barker WC, Boeckmann B, et al. The Universal Protein Resource (UniProt): an expanding universe of protein information. Nucleic Acids Res. 2006;34(Database issue):D187–91.

22. Efremova M, Vento-Tormo M, Teichmann SA, Vento-Tormo R. CellPhoneDB: inferring cell-cell communication from combined expression of multi-subunit ligand-receptor complexes. Nat Protoc. 2020;15(4):1484–506.

23. Vento-Tormo R, Efremova M, Botting RA, Turco MY, Vento-Tormo M, Meyer KB, et al. Single-cell reconstruction of the early maternal-fetal interface in humans. Nature. 2018;563(7731):347–53.

24. Page L, Brin, S., Motwani, R. and Winograd, T.,. The PageRank Citation Ranking: Bringing Order to the Web. Stanford InfoLab. 1999.

25. Kim Y, Choi S. Nonnegative tucker decomposition. 2007 IEEE Conference on Computer Vision and Pattern Recognition. 2007:1–8.

26. Zhou JX, Taramelli R, Pedrini E, Knijnenburg T, Huang S. Extracting intercellular signaling network of cancer tissues using ligand-receptor expression patterns from whole-tumor and single-cell transcriptomes. 2017.

27. Aghdashi MA, Seyedmardani S, Ghasemi S, Khodamoradi Z. Evaluation of Serum Calprotectin Level and Disease Activity in Patients with Rheumatoid Arthritis. Curr Rheumatol Rev. 2019;15(4):316–20.

28. Wang Q, Chen W, Lin J. The Role of Calprotectin in Rheumatoid Arthritis. J Transl Int Med. 2019;7(4):126–31.

29. Wu FP, Gao JF, Kang J, Wang XX, Niu Q, Liu JX, et al. B Cells in Rheumatoid Arthritis:Pathogenic Mechanisms and Treatment Prospects. Frontiers in Immunology. 2021;12.

30. Dienz O, Rincon M. The effects of IL-6 on CD4 T cell responses. Clin Immunol. 2009;130(1):27–33.

31. Tirosh I, Izar B, Prakadan SM, Wadsworth MH, 2nd, Treacy D, Trombetta JJ, et al. Dissecting the multicellular ecosystem of metastatic melanoma by single-cell RNA-seq. Science. 2016;352(6282):189–96.

32. Johnstone CN, Chand A, Putoczki TL, Ernst M. Emerging roles for IL-11 signaling in cancer development and progression: Focus on breast cancer. Cytokine Growth Factor Rev. 2015;26(5):489–98.

33. Liao M, Liu Y, Yuan J, Wen Y, Xu G, Zhao J, et al. Single-cell landscape of bronchoalveolar immune cells in patients with COVID-19. Nat Med. 2020;26(6):842–4.

34. Guo C, Li B, Ma H, Wang X, Cai P, Yu Q, et al. Single-cell analysis of two severe COVID-19 patients reveals a monocyte-associated and tocilizumab-responding cytokine storm. Nat Commun. 2020;11(1):3924.

35. Højen JF, Kristensen MLV, McKee AS, Wade MT, Azam T, Lunding LP, et al. IL-1R3 blockade broadly attenuates the functions of six members of the IL-1 family, revealing their contribution to models of disease. Nature immunology. 2019;20(9):1138–49.

36. Butler D, Mozsary C, Meydan C, Foox J, Rosiene J, Shaiber A, et al. Shotgun transcriptome, spatial omics, and isothermal profiling of SARS-CoV-2 infection reveals unique host responses, viral diversification, and drug interactions. Nature communications. 2021;12(1):1–17.

37. Weinreb C, Wolock S, Klein AM. SPRING: a kinetic interface for visualizing high dimensional single-cell expression data. Bioinformatics. 2018;34(7):1246–8.

38. Chamberlain M, Hanamsagar R, Nestle FO, de Rinaldis E, Savova V. Cell type classification and discovery across diseases, technologies and tissues reveals conserved gene signatures and enables standardized single-cell readouts. bioRxiv. 2021.

39. Korsunsky I, Millard N, Fan J, Slowikowski K, Zhang F, Wei K, et al. Fast, sensitive and accurate integration of single-cell data with Harmony. Nat Methods. 2019;16(12):1289–96.

